# Temporal resolution of *C. vicina* compound eyes and ocelli

**DOI:** 10.64898/2026.10.15.759508

**Authors:** Grace C. Hickey, Marisa S. McDonald, Daniel R. Chappell, Martin F. Wehling

## Abstract

Most flying insects have two visual systems which serve different purposes depending on the insect. The compound eyes are the main visual system and provide the insect with better spatial vision, while the ocelli act as a secondary visual system and often provide faster processing at a lower spatial resolution. To accomplish this, ocelli can have widely varying characteristics, including temporal resolution. The goal of this study was to examine the critical flicker fusion frequency (CFF) of the blowfly *Calliphora vicina*’s compound eyes and ocelli using electroretinography. We found that *C. vicina*’s compound eyes and ocelli both have a maximum CFF of 240–250 Hz, which is consistent with its natural light environment, flight behaviors, and the CFF values found for other Dipteran species.

**Summary:** This study examined the flicker fusion frequency of the blowfly *Calliphora vicina* using electroretinography. We found a CFF of ∼250 Hz, consistent with their environment, behaviors, and other Dipteran species.

## Introduction

Visual systems are used throughout the animal kingdom to perform critical ecological tasks, including navigation, orientation, mate recognition, hunting, and more. Insects typically have two main compound eyes composing the primary visual system, but many insects also have a secondary visual system known as the ocellar system. While there are only a few examples of walking insects with ocelli, almost all flying insects have ocelli, and their functions appear to be broadly flight-related (Krapp, 2009).

The ocellar system is typically composed of three simple eyes in a triangular formation on the surface of the head between the main compound eyes. A common feature in many ocelli, though not all, is a foveal streak which appears to align with the horizon while the insect is in flight to provide higher spatial resolution in this area (for review, Ribi and Zeil, 2018; in dragonflies, Stange et al., 2002). Dipteran ocelli are generally thought to be under-focused, with the lens focusing images well behind the retina, indicating that they are not image forming (e.g. Cornwell, 1955; Schuppe and Hengstenberg, 1993), but may still distinguish coarse patterns (Schuppe and Hengstenberg, 1993). Because of the rapid processing and under-focused lens, Dipteran ocelli may not be used for image formation or spatial resolution, but instead to assist with specific tasks, such as enhancing flight stability by tracking changes in light intensities across each ocellus (Krapp, 2009; Parsons et al., 2006).

Depending on the needs of the insect, ocelli are hypothesized to have a wide range of specific functions and display structural changes which reflect those functions. There is wide variation in the sensitivity, speed, and neural processing pathways of ocellar systems which broadly correspond to insects’ typical light environment (Mizunami, 1995). This can be seen even in closely related species, as ants with different temporal niches of the same genus, as well as walking vs. flying individuals of the same species, have different ocellar sizes and structures which reflect their ecological niche (Narendra and Ribi, 2017).

A visual system must have sufficient temporal resolution to keep up with its animal’s movement and environment. One measure of temporal resolution is critical flicker fusion frequency (CFF), which reflects the frequency of a flashing stimulus at which an eye can no longer distinguish individual flashes from a steady light. For humans, CFF falls around 50-90 Hz (Mankowska et al., 2021), while other species can have very different CFFs depending on their visual systems, typical light environments, and ecological niches. For example, while the CFF of aquatic animals tends to be significantly slower than terrestrial animals, the CFF of the snapping shrimp *Alpheus heterochaelis* is at least 160 Hz, which is attributed to keeping up with its own rapid fire snapping movements (Kingston et al., 2020).

Flying animals—generally birds and insects—have faster CFFs than non-flying animals (Inger et al., 2014; Lafitte et al., 2022). The average flicker fusion frequency of diurnal bird species falls around 90 Hz (Inger et al., 2014), with predatory birds such as the peregrine falcon ranging up to 125 Hz (Potier et al., 2020). Insects exhibit a massive range of flicker fusion frequencies. Nocturnal and slower-moving insects tend to have slow CFFs, averaging around 80 Hz (Inger et al., 2014) and with some well under this mark (e.g. *Periplaneta americana* at approximately 40 Hz; Miall, 1978). Diurnal, fast-flying species of insects tend to have the fastest CFFs, many with a flicker fusion frequency well above 200 hertz for the compound eye (Autrum and Hoffman, 1960; Inger et al., 2014; Miall, 1978; Ruck, 1958; Ruck, 1961). The compound eye of *Calliphora vicina* was previously found to have a CFF of at least 265 Hz (Autrum, 1950).

Ocellar systems have faster neural processing speeds than compound eyes. This is because there are fewer interneurons which separate the ocelli from the optic lobe than the compound eyes, and the associated interneurons have larger diameters and thus higher propagation speeds (Jean-Guillaume and Kumar, 2022; Mizunami, 1995). However, CFF studies on insects are still relatively sparse (Inger et al., 2014; Lafitte et al., 2022); the CFFs of only around 150 species have been studied, with very few of these being insects and an even smaller subset focusing on insect ocellar systems. The current study is one of few which has examined the electroretinographic properties of ocelli, especially flicker fusion frequency, in any insect species.

In this study, we examined the Dipteran blowfly, *Calliphora vicina. C. vicina* has long been studied to better understand the functional role of visual systems in flight (Cornwell, 1955; Parsons et al., 2006; Parsons et al., 2010; Schuppe and Hengstenberg, 1993), and the neural structure of their visual systems is well documented (e.g. Nässel and Hagberg, 1985; Simmons et al., 1994; Strausfeld and Bassemir, 1985). From previous work, we know that *Calliphora* ocelli produce under-focused images, indicating they are likely not being used for spatial resolution, but for other visual tasks (Cornwell, 1955; Schuppe and Hengstenberg, 1993). Behavioral evidence has previously suggested that one role of ocelli in *C. vicina* is controlling light responses, including the dorsal light response, or the ability of an insect to orient their dorsal side towards light (Schuppe and Hengstenberg, 1993), or by assisting in positive phototactic movements in conjunction with the compound eyes (Cornwell, 1955). In *C. vicina*, most early electrophysiological work focused on the spectral sensitivity and field of view of the ocelli (Kirschfeld and Lutz, 1977; Schuppe and Hengstenberg, 1993), but ocellar CFF has never been examined in *C. vicina*.

This research will inform our understanding of ocellar function, including how insects may be affected by artificial light, and will improve the potential for bio-inspired design based on ocellar systems. We used electroretinography to determine maximum critical flicker fusion frequency of adult *C. vicina* compound eyes and ocelli under dark and white light-adapted conditions, as temporal resolution tends to increase under light-adapted conditions (Laughlin and Weckström, 1993). Based on their flight behaviors (Bomphrey et al., 2009), typical bright-light environments (Arias-Robledo et al., 2019), and the flicker fusion frequencies of other Dipteran species (Autrum, 1950; Autrum and Hoffman, 1960; Miall, 1978; Ruck, 1958; Ruck, 1961), we expect that *C. vicina* compound eyes and ocelli will both have relatively high CFFs compared to other insects. Additionally, we hypothesized the ocelli would have higher CFFs than compound eyes to assist in the rapid information collection and processing required by flying insects, and which ocelli are hypothesized to play a role in.

## Methods

### Animals

In this study, we examined the blowfly *Calliphora vicina* (formerly *Calliphora erythrocephala*). The flies were ordered as maggots (Amazon), then pupated and eclosed in the laboratory. As adults, they had constant access to water, sugar water, and beef liver. We replenished water and sugar daily, and replaced liver weekly. Flies were housed in mesh cages next to a window with natural lighting, thus experiencing the natural dark/light cycle for northern Florida in the summer (approximately 10 hours dark:14 hours light). Room temperature was maintained at approximately 22º–24ºC. We used all female flies, as male ocelli were too obstructed by the compound eyes and face bristles to consistently place an electrode, and we only tested flies once they were fully sclerotized to ensure effective electrode placement.

### Electrophysiology

#### Electroretinography Setup

We used single-ended, extracellular electroretinogram (ERG) recordings in this study. During experimentation, we immobilized flies with wax on a custom insect holder and mounted them in a light-tight Faraday cage. We used electrolytically sharpened tungsten electrodes and placed the recording electrode subcorneally in the compound eye or ocellus with a ground electrode in the contralateral compound eye.

We used a custom electroretinography setup for all electrophysiology experiments. The light stimulus was a xenon arc lamp (Newport Model 66905 housing, Newport Model 6255 150W lamp), which passed through a pair of rotating gradient neutral density filters (Thorlabs model NDC-100C-4M, Newport Motion control model ESP301) to control light intensity. The flickering light stimulus was created using an optical chopper (Thorlabs MC2000B), and the light stimulus was then focused onto a UV-VIS liquid light guide (Newport 77569) that terminated at the eye. During experimentation, AC signals were amplified with a high-pass frequency of 0.3Hz and a low-pass frequency of 10kHz (NPI DPA-2FS) and recorded with a data acquisition system (National Instruments PCI-7221with BNC-2090A terminal block). We used a custom LabView program to control all components, as well as to view and digitize all recordings which were stored for later analysis.

#### Experimentation

We measured the CFF_max_, or the temporal rate at which the eye is unable to follow increasing rates of stimulus flashes even with increasing irradiance. We tested insects at three levels of increasing light intensity (1x10^19^, 1x10^20^, and 1x10^21^ photons cm^-2^ sec^-1^), under both dark and white light adaptation, to determine the CFF_max_ of their visual system. These irradiance levels are brighter than broad daylight, ensuring we reach the CFF_max_. For both adaptive states, we adapted all animals for a minimum of 20 minutes, or until there was a consistent response to a test flash of light. During the white light adaptation, we used a secondary light source (Ocean Optics HL-2000) and fiber optic cable (Ocean Optics P1000-2-UV-VIS) to focus white light onto the retina at an intensity of approximately 1x10^16^ photons cm^-2^ sec^-1^. The adapting light remained on throughout experimentation. During experimentation, we presented animals with 20 cycles of a flickering white light stimulus from 50-300 Hz at 10 Hz steps with rest periods of 60 s between trials.

#### Statistics

To assess the range of frequencies at which the compound eye and ocelli were able to follow a flickering light stimulus, we prepared the electrophysiological recordings for analysis in a custom R program. The beginning and end 10% of each recording was removed to decrease anomalies in the data caused by eye on/off dynamics. We detrended the data using a cubic spline fit to the raw electrophysiological recordings. Example detrended flicker traces from *C. vicina* compound eyes are given in Fig. 2D,H. Next, we ran a Fourier transform on the detrended data to determine the relative signal power for the trial at each frequency of stimulation and generated a relative power versus stimulus frequency curve for each eye. The power curves were normalized and averaged for each of the trial sets. A 5% relative power threshold was used to determine if the eye was following the flickering light stimulus at a particular frequency. To determine the CFF_max_, we considered only light-adapted trials at the irradiance for which CFF was the highest. The CFF_max_ given here represents the frequency at which the lower bound of the 95% confidence interval of the averaged relative powers of the FFTs crossed the 5% threshold. We discarded trials which showed significant noise in the raw electrophysiology trace.

**Figure 1.**
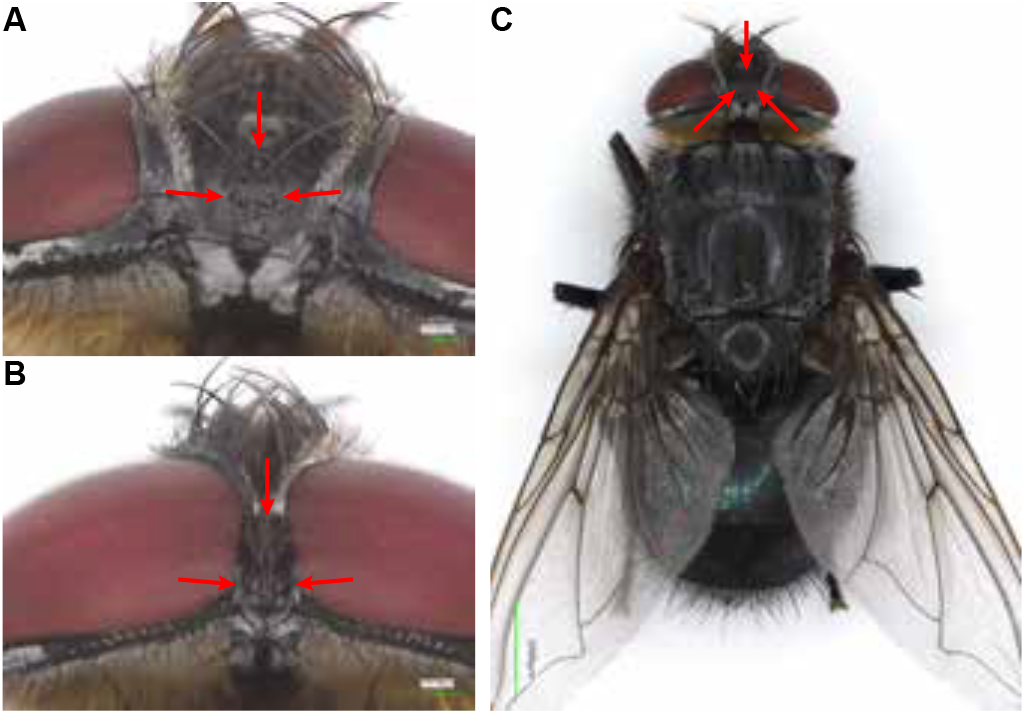
Calliphora vicina eyes and body. Red arrows indicate the position of the ocelli. A) The head of a female *C. vicina* (100x). B) The head of a male *C. vicina* (100x). Note that the median ocellus is obstructed in this view. C) The body of a female *C. vicina* (20x).

**Figure 2.**
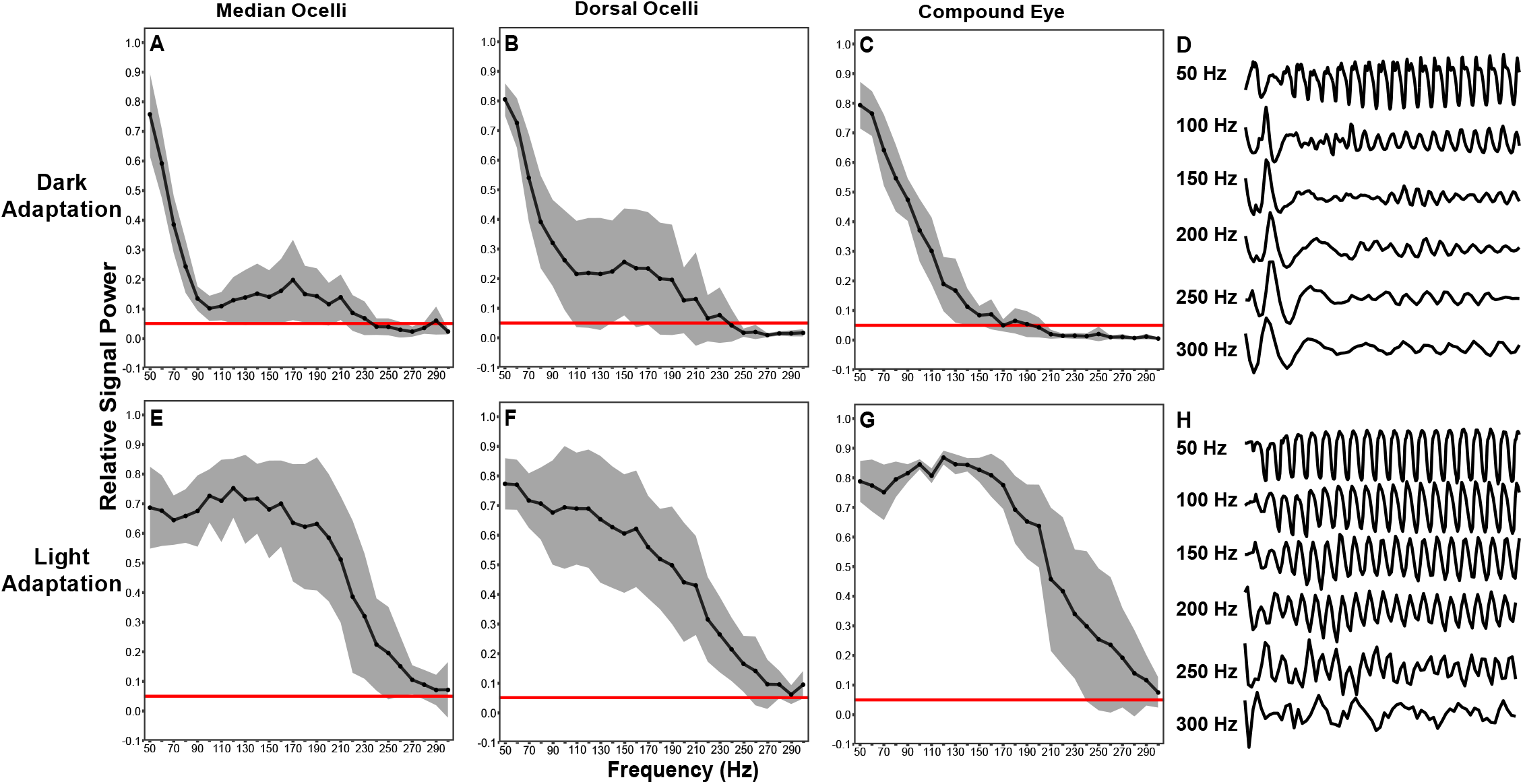
Results of FFT analysis and sample flicker traces. (A-C, E-F) The gray ribbon indicates the 95% confidence interval, while the line represents the mean signal power at each frequency for the given eye. The maximum CFF listed reflects the frequency at which the lower end of the 95% confidence interval crosses the 5% relative power threshold, shown by the red line. For all light adapted trials, the means and upper end of the confidence intervals remain above 300 Hz, indicating that the maximum CFF of some individuals may be well above 300 Hz. A) Dark-adapted median ocelli. CFF_max_ of at least 210 Hz. n=8. B) Dark-adapted dorsal ocelli. CFF_max_ of at least 150 Hz. n=8. C) Dark-adapted compound eye. CFF_max_ of at least 140 Hz. n=7. E) Light-adapted median ocelli. CFF_max_ of at least 240 Hz. n=6. F) Light-adapted dorsal ocelli. CFF_max_ of at least 250 Hz. n=8. G) Light-adapted compound eye. CFF_max_ of at least 240 Hz. n=6. (D, H) Sample flicker traces from *C. vicina* compound eyes. D) FFT analysis indicated that the eye followed the stimulus at 50, 100, and 150 Hz, but not at 200, 250, or 300 Hz, indicating that we reached the critical flicker fusion frequency. H) FFT analysis indicated that the eye followed the stimulus at all tested frequencies, indicating that we did not reach the critical flicker fusion frequency for this individual.

## Results and Discussion

The CFF values were obtained for each eye (compound eyes, median ocelli, and dorsal ocelli), irradiance, and light adaptation (Figs 2A-C, 2E-G). Within each eye, mean CFF value did not vary significantly between irradiance levels, indicating that we reached the maximum critical flicker fusion frequency (CFF_max_) (as described by Frank, 1999). We found that the CFF_max_ of light-adapted *C. vicina* compound eyes is at least 240 Hz. The CFF_max_ for the medial ocelli was at least 240 Hz, and at least 250 Hz for dorsal ocelli. The upper bound of the 95% confidence interval of the averaged relative powers of the FFTs extended past 300 Hz for all three eyes, indicating that the maximum CFFs of some individuals may be beyond the tested frequencies. There were no significant differences between any of the eyes’ CFF_max_ values. The results of our FFT analysis are shown in Fig. 2. The measured CFF values agree with the CFF values obtained in temporal resolution studies on other Dipteran species, for both compound eyes and ocelli (e.g. Eichorn et al., 2017; Mebourou et al., 2018; Miall, 1978; Ruck, 1958), including one previously measured value for *C. vicina*’s compound eye (265 Hz in Autrum, 1950).

The average CFF for light-adapted eyes was clearly higher than dark-adapted eyes, though statistical significance varied. Light-adapted trials had CFFs up to or over 100 Hz higher than their dark-adapted counterparts (Fig. 2). This phenomenon has been seen in studies of other arthropods from crustaceans to insects (Frank, 2003; Laughlin and Weckström, 1993; Miall, 1978). Additionally, the ‘fast cells’ found in the eyes of other highly maneuverable, diurnal Dipteran species have significantly increased temporal resolution under light-adapted conditions compared to dark-adapted conditions, consistent with what is seen in *C. vicina* (Laughlin and Weckström, 1993).

Similarly, the temporal resolution of an animal is often linked to its natural light environment, as eyes must trade light sensitivity for temporal resolution (Inger et al., 2014); thus, diurnal insect species tend to have significantly higher CFFs than nocturnal species (Laughlin and Weckström, 1993). More generally, insects typically exposed to high and variable light levels in their natural environment have a higher CFF (mean 167 Hz, Lafitte et al., 2022) than insects exposed to low light levels (mean 77 Hz, Lafitte et al., 2022). The high flicker fusion frequencies found in this study are consistent with the typical environments of *C. vicina*, which include both bright, open field environments and more shaded wooded areas (Arias-Robledo et al., 2019). There is a similar trend in another Calliphorid fly, *L. sericata*, which has a CFF_max_ of over 200 Hz (Ruck, 1961) and is also found in open, bright environments (Arias-Robledo et al., 2019).

Additionally, one could expect flicker fusion frequency to increase as animal speed increases. This trend has been observed in aquatic animals, such as the snapping shrimp *A. heterochaelis*, which is fast-moving and must react quickly in its environment, and has the highest known maximum CFF of any crustacean or aquatic animal (Kingston et al., 2020). Flying animals also tend to have the fastest flicker fusion frequencies, regardless of taxa (Inger et al., 2014; Lafitte et al., 2022). *C. vicina* have a moderate flight speed for Dipteran species, with the most common flight speeds lying between 1.2 m/s and 2.3 m/s during normal flight (Bomphrey et al., 2009). This is slightly faster than the normal flight speeds of *Drosophila*, which are around 1 m/s and accompanied by a lower CFF of 50-80 Hz (Lafitte et al., 2022; Leitch et al., 2021). Calyptrate flies, a subsection of Diptera which includes families like Calliphoridae, Muscidae, and Glossinidae, tend to have fast take off times that are mediated at least in part by the halteres (Yarger et al., 2021), and it is possible that they have also developed faster vision to help process the visual information encountered during this fast take off.

While insects’ temporal resolution may be tuned to their natural light environment, increases in artificial light have created another light environment for insects to contend with. Because of their high CFF_max_, insects such as *C. vicina* are likely to perceive the flicker of artificial lighting, such as fluorescent lights which flicker around 100-120 Hz. In nature, artificial lighting can have a huge range of effects on insects, such as negative impacts on foraging behaviors, reproduction, development, and increased vulnerability to predation (Owens et al., 2020). The flicker of artificial light at night may be particularly detrimental to species with CFFs high enough to see it (Lafitte et al., 2022). Understanding the CFF of more species will help understand and mitigate the effects of flicker in artificial lighting on insects, for example by informing appropriate use of artificial light sources, especially outdoors, where it may negatively impact insect populations. Given the important ecological roles of insects, understanding how artificial light environments may impact their behavior is necessary.

The current study is one of few which has examined the electroretinographic properties of ocelli, especially flicker fusion frequency. Future work in this area should include new and updated research into the CFFs of various insect species, as well as characterizing the light environments and behavioral characteristics of more species to determine CFF’s role in behavior and ecological adaptation. Additionally, anatomical and electrophysiological studies of the downstream neural processing and integration of visual information from compound eyes and ocelli will further clarify the functions of ocelli. This research will be important across fields including bio-inspired design and visual ecology, helping clarify the diverse roles of ocelli and how we can mimic their functionality.

## Acknowledgements

We would like to acknowledge the Universities Space Research Association’s AFRL Summer Scholars Program for providing the opportunity to conduct this research.

## Competing Interests

The authors declare no competing or financial interests.

## Author Contributions

Conceptualization: G.C.H., M.S.M.

Methodology: G.C.H., M.S.M.

Software: D.R.C.

Formal analysis: G.C.H.

Investigation: G.C.H., M.S.M.

Resources: M. W.

Data curation: G.C.H., D.R.C.

Writing – original draft preparation: G.C.H

Writing – review and editing: G.C.H., M.S.M, D.R.C., M. W.

Visualization: G.C.H.

Funding acquisition: M.W.

## Funding

Funding for this project was provided by the AFRL Air Warfare Directorate and Universities Space Research Association through the AFRL Summer Scholars internship program. This work is available for public release under public release number: AFRL-2025-4436.

## Data Availability

Data and code are available upon request.

## Notes

### Competing Interest Statement

The authors have declared no competing interest.

